# Unraveling Spatial Gene Associations with SEAGAL: a Python Package for Spatial Transcriptomics Data Analysis and Visualization

**DOI:** 10.1101/2023.02.13.528331

**Authors:** Linhua Wang, Chaozhong Liu, Zhandong Liu

## Abstract

**Summary:** In the era where transcriptome profiling moves towards single-cell and spatial resolutions, the traditional co-expression analysis lacks the power to fully utilize such rich information to unravel spatial gene associations. Here we present a Python package called Spatial Enrichment Analysis of Gene Associations using L-index (SEAGAL) to detect and visualize spatial gene correlations at both single-gene and gene-set levels. Our package takes spatial transcriptomics data sets with gene expression and the aligned spatial coordinates as input. It allows for analyzing and visualizing spatial correlations at both single-gene and gene-set levels. The output could be visualized as volcano plots and heatmaps with a few lines of code, thus providing an easy-yet-comprehensive tool for mining spatial gene associations.

**Availability and Implementation:** The Python package SEAGAL can be installed using pip: https://pypi.org/project/seagal/. The source code and step-by-step tutorials are available at: https://github.com/linhuawang/SEAGAL.

## 1 Introduction

Spatial Transcriptomics (ST) is a molecular profiling technique that maps gene expression across a tissue sample at the single-cell level, offering a comprehensive overview of gene expression patterns and unique biological insights (Marx, 2021). The technique has applications in various fields, such as developmental biology (Choe, 2023), neuroscience (Close, 2021), and cancer research (Anderson, 2022).

To aid in data preprocessing and analysis, a variety of computational tools have been developed. Among these are SpatialDE (Svensson, 2018) and SquidPy (Palla, 2022), which identify Spatial Variable Genes (SVGs) through variance decomposition or spatial autocorrelation. These tools bring the concept of Highly Variable Genes (HVGs) from single-cell analysis to spatial transcriptomics. However, these SVG tools do not account for spatial associations such as colocalization and exclusion of paired features (genes or gene groups). Such spatial associations would add another dimension to traditional correlation-based co-expression analysis methods like the Weighted Correlation Network Analysis (Langfelder, 2008).

To fill the gap, we developed SEAGAL for Spatial Enrichment Analysis of Gene Associations using L-index, a bivariate spatial association measure proposed by (Lee, S.-I., 2001). Based on the L-index, SEAGAL allows ST data preprocessing, SVG detection, Spatially Associated Genes (SAG) identification, immune colocalization pattern recognition in local tissue niches, SAG-based gene module discovery, and so on.

SEAGAL is written in Python and compatible with well-established single-cell and spatial-omics analytical packages such as Scanpy (Wolf, 2018) and SquidPy (Palla, 2022). Moreover, integrated with well-supported Python visualization packages, including matplotlib and seaborn, SEAGAL allows visualizing each data analysis result in spatial heat maps, violin plots, and cluster maps. Its major functionalities could be carried out in a few lines of code by following the user manual and tutorials available at GitHub: https://github.com/linhuawang/SEAGAL.

## 2 Package description

SEAGAL takes inputs from either of the two types: (1) Raw 10X Visium data output from SpaceRanger, and (2) a user-processed folder containing CSV-format files, including a raw count matrix, and metadata with x and y coordinates as columns. After importing the module `seagal`, users can load either type of input using the command `load_raw()`.

### Preprocess

As a quality control step, SEAGAL removes genes observed in less than ten spots and spots with less than 150 UMI counts. To adjust for library size and normalize the variance, it transforms the raw counts into log-scaled count-per-million data types. Both steps are completed by calling the function `process_st()`.

### Immune-colocalization

`group_adata_by_genes(grouped=True)` followed by `spatial_association()` allow users to group gene markers to identify cell types’ spatial associations, such as immune colocalization investigation. Users could either use default immune markers provided by SEAGAL or provide their marker dictionary as an optional parameter.

### Gene-gene spatial association

To detect gene-gene spatial associations, users must first call `spatial_pattern_genes()` to detect SVGs. We recommend using <1000 top SVGs or SVGs with Moran’s I > 0.3 by specifying parameters `I` or `topK`. Then, `group_adata_by_genes(grouped=False)`will yield the spatial gene association for the selected top SVG pairs.

### Gene module detection

Function `genemodules()` allows automatic gene module identification by finding the number of clusters that maximizes the silhouette score in iterative hierarchical clustering runs.

### Visualization

- General spatial associations could be visualized in a volcano plot and facilitate users to select their SAGs using `volcano()`.
- Spatial heat map of gene-gene association or immune-colocalization for paired variables could be visualized by `hotspot()`.
- Summarized immune colocalization patterns could be visualized in a clustered heat map via `clustermap()`.
- Gene modules’ expression patterns could be visualized through `module_pattern()`. And `module_hotspot()` plots module-module associations in spatial heat maps.

## 3 Example

In this section, we will use a 10X Visium dataset as an example to explore SEAGAL’s various functionalities. Other tutorials including how to use CSV-format input or how to use user-defined marker gene lists for exploring cell-cell colocalization and exclusion are available at: https://github.com/linhuawang/SEAGAL.

The Human Breast Cancer block 1 sample (HBC1) was originated from 10X Genomics and could also be downloaded from our example datasets: https://github.com/linhuawang/SEAGAL.

Figure 1 shows the plots generated by executing the code below using the sample data.

**Fig. 1.**
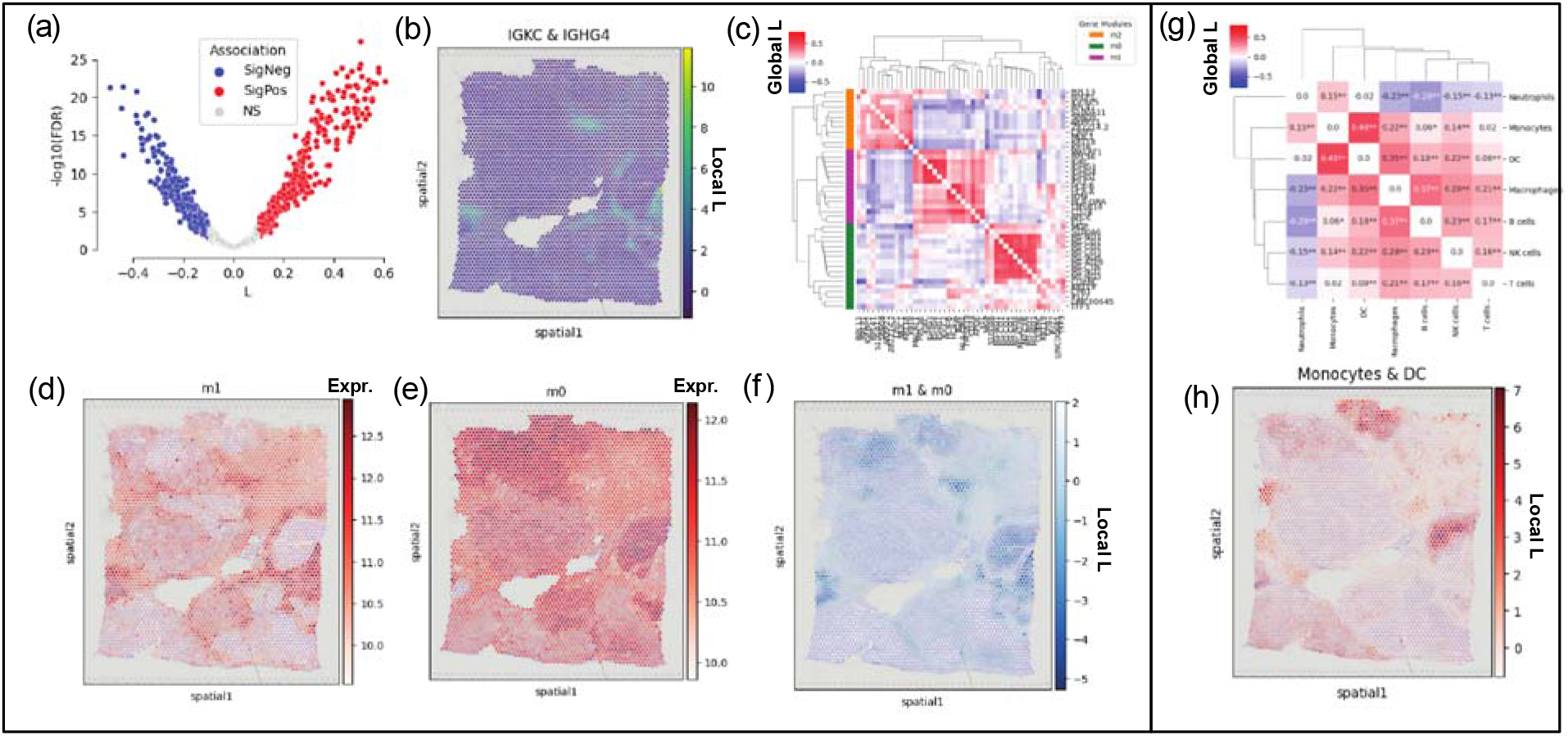
**(a)** Volcano plot of Spatially Associated Gene (SAG) pairs; each dot is a pair of spatially variable genes; x axis shows the global spatial association in L-index and y axis shows the -log10 of false discovery rate in permutation test. The significance is colored red (significantly positive), blue (significantly negative) and gray (insignificant). **(b)** Spatial heat map showing niches where a pair of SAGs are colocalized. **(c)** Clustered heatmap with gene modules colored in each row. **(d-e)** Detected modules’ expression patterns in spatial heat maps. **(f)** Spatial heat map showing local niches where a pair of modules expression negatively associated. **(g)** Clustered heatmap for immune cell types’ spatial associations. **(h)** Spatial heat map showing the spatial niches where Monocytes and Dendritic Cells (DC) are highly colocalized.

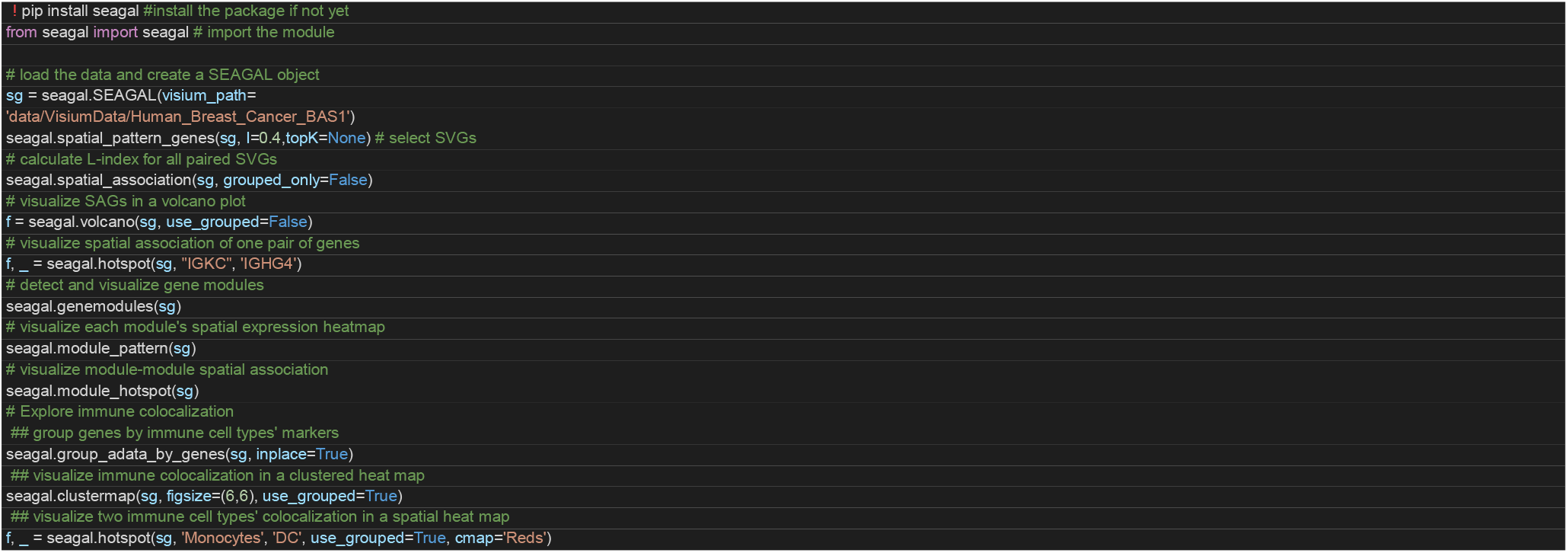

## Acknowledgements

We thank Professor Xiang H.-F. Zhang and Dr. Yang Gao for initial discussions on the gap in studying spatial variable associations of immune cells that motivated us to develop SEAGAL.

## References

Anderson, A. C. et al. (2022) Spatial transcriptomics. Cancer Cell, 40(9), 895–900.

Choe, K. et al. (2023) Advances and Challenges in Spatial Transcriptomics for Developmental Biology. Biomolecules, 13(1), 156.

Close, J. L. et al. (2021) Spatially resolved transcriptomics in neuroscience. Nature Methods, 18(1), 23–25.

Langfelder, P. and Horvath, S. (2008) WGCNA: an R package for weighted correlation network analysis. BMC Bioinformatics, 9, 559.

Lee, S.-I. (2001) Developing a bivariate spatial association measure: An integration of Pearson’s r and Moran’s I. Journal of Geographical Systems, 3, 369–385.

Marx, V. (2021) Method of the Year: spatially resolved transcriptomics. Nature Methods, 18(1), 9–14.

Palla, G. et al. (2022) Squidpy: a scalable framework for spatial omics analysis. Nature Methods, 19, 171–178.

Svensson, V. et al. (2018) SpatialDE: identification of spatially variable genes. Nature methods, 15, 343–346

Wolf, F. A. et al. (2018) SCANPY: large-scale single-cell gene expression data analysis. Genome Biology, 19, 15.

